# Improved Placement of Multi-Mapping Small RNAs

**DOI:** 10.1101/044099

**Authors:** Nathan R. Johnson, Jonathan M. Yeoh, Ceyda Coruh, Michael J. Axtell

**Author notes:** Current address: Salk Institute for Biological Studies, La Jolla, CA, 92037 USA. Author for Correspondence: Michael J. Axtell Department of Biology, 208 Mueller lab, 16802.

## Abstract

High-throughput sequencing of small RNAs (sRNA-seq) is a popular method used to discover and annotate microRNAs (miRNAs), endogenous short interfering RNAs (siRNAs) and Piwi-associated RNAs (piRNAs). One of the key steps in sRNA-seq data analysis is alignment to a reference genome. sRNA-seq libraries often have a high proportion of reads which align to multiple genomic locations, which makes determining their true origins difficult. Commonly used sRNA-seq alignment methods result in either very low precision (choosing an alignment at random) or sensitivity (ignoring multi-mapping reads). Here, we describe and test an sRNA-seq alignment strategy that uses local genomic context to guide decisions on proper placements of multi-mapped sRNA-seq reads. Tests using simulated sRNA-seq data demonstrated that this local-weighting method outperforms other alignment strategies using three different plant genomes. Experimental analyses with real sRNA-seq data also indicate superior performance of local-weighting methods for both plant miRNAs and heterochromatic siRNAs. The local-weighting methods we have developed are implemented as part of the sRNA-seq analysis program ShortStack, which is freely available under a general public license. Improved genome alignments of sRNA-seq data should increase the quality of downstream analyses and genome annotation efforts.

## Introduction

High-throughput sampling of cDNA libraries derived from endogenous small RNAs (sRNA-seq) is a proven and widely used method. sRNA-seq allows both discovery and quantification of diverse regulatory small RNAs, which can include microRNAs (miRNAs), endogenous short interfering RNAs (siRNAs) and Piwi-associated RNAs (piRNAs), depending on the species and specimen. sRNA-seq data have played critical roles in the advancement of understanding of miRNAs, siRNAs, and piRNAs from numerous species. However, despite the large amounts of available sRNA-seq data, multiple issues remain. For example, the central miRNA database, miRBase, is thought to contain substantial numbers of incorrect annotations (Kozomara and Griffiths-Jones 2014; Taylor *et al*. 2014). Additionally, annotation of siRNA loci is much less well developed, especially in plants where siRNAs frequently represent the majority of expressed small RNAs (Coruh *et al*. 2014). Many factors contribute to annotation errors and omissions, including the use of sub-optimal methodologies for small RNA-seq alignments to reference genomes.

Alignment of small RNA-seq data to a reference genome remains a persistent, if perhaps under-recognized, problem. A major issue is the prevalence of multi-mapping (MMAP) reads in sRNA-seq data. MMAP reads occur when there are multiple best-scoring alignments to the reference genome. MMAP reads are quite rare in modern polyA+ mRNA-seq data due to their longer read-lengths, and due to the fact that polyA+ mRNAs generally are transcribed from single-copy sequences. In contrast, MMAP reads are much more frequent in sRNA-seq data due both to the short lengths of the reads and their tendency to originate from higher-copy number regions of the genome. Endogenous siRNAs are known to come from repetitive regions of many genomes (Matzke and Mosher 2014), while identical miRNAs are often encoded by multiple paralogous loci (Cuperus *et al*. 2011).

MMAP sRNA-seq reads are often dealt with simplistically, either by randomly selecting one the possible alignment positions, or by ignoring them entirely. For instance, the popular bowtie aligner (Langmead et al. 2009) by default randomly selects one position for MMAP reads, and can also be configured to ignore them. Both of these approaches have the advantage of computational speed, but both also have significant downsides: random selection results in large error rates, while ignoring MMAP reads discards large portions of sRNA-seq libraries. More sophisticated approaches for placing MMAP reads have been described for mRNA-seq data. Expression estimation using the ERANGE method, where the expression of MMAP reads is measured as a proportion of uniquely mapped reads within a particular read-cluster, was shown to improve estimates of mRNA abundance from mRNA-seq data (Mortazavi *et al*. 2008). This approach has been applied to sRNA-seq with the SiLoCO method (Moxon *et al*. 2008; Stocks *et al*. 2012), where loci are identified by clustering and MMAP reads lend abundance proportional to their MMAP-value. A similar approach applied to cap analysis of gene expression (CAGE) data instead weights only by the number of different species of uniquely aligned reads, preventing highly expressed sequences from lending proportionally higher weight (Faulkner *et al*. 2008). Both of these methods improve the accuracy of mRNA quantification. The Rcount method uses similar ideas (Schmid and Grossniklaus 2015) but produces an alignment output, as opposed to solely a quantification of mRNA expression levels.

sRNA-seq and mRNA-seq data are similar in that both data types are expected to frequently result in local genomic clusters of aligned reads with distinct sequences. For mRNA-seq, clustering of alignment positions results from experimentally induced fragmentation of longer mRNAs in preparation for cDNA synthesis and sequencing. In sRNA-seq, nature performs the fragmentation through various RNA processing events acting on longer precursor RNAs. For miRNAs, the precursor stem-loop RNA often produces multiple variants of the major miRNA, including miRNA*'s and isomirs (Coruh *et al*. 2014). Endogenous siRNAs from both animals and plants are also often produced from longer hairpin or dsRNA precursors that spawn multiple distinct siRNAs (Allen *et al*. 2005; Okamura *et al*. 2008), or from short dsRNA precursors that are themselves spawned from co-located genomic clusters (Blevins *et al*. 2015; Zhai *et al*. 2015). Finally, piRNAs are also produced in large genomic clusters in multiple animals (Aravin *et al*. 2006; Malone *et al*. 2009). We thus reasoned that the biologically expected clustering of sRNA-seq reads at their true loci of origin would enable an ERANGE-or Rcount-like algorithm to improve placement of MMAP sRNA-seq reads. All of the previously described implementations of this general read-rescue strategy have specific features and settings particular to mRNAs and hence are not directly useful for sRNA-seq analysis. Thus, we implemented these general ideas into the alignments performed by an updated version of our previously described sRNA-seq analysis tool, ShortStack (Axtell 2013). Here, we describe the implementation and testing of sRNA-seq alignment performance using a local-weighting method to better place MMAP small RNA reads.

## Materials and Methods

### Alignment methods

In thinking about treatment of MMAP sRNA-seq reads, we make two assumptions. First, we assume that each read in a sRNA-seq experiment represents a single small RNA molecule, which therefore must have had a single genomic origin. Note that this assumption does not mean we assume that all reads with the same sequence necessarily have the same genomic origin. For instance, if we find 100 reads of identical sequence with a MMAP value of two (e.g., two possible alignment positions), it's certainly possible that some of those reads came from one location, and the rest from the other. In other words, we treat single reads as indivisible, but multiple reads with the same sequence can each be placed in a different location. Second, we assume that the goal of sRNA-seq alignment to the reference genome is to identify the site of transcriptional origin of the small RNAs, not to list their possible targets. Target predictions, which usually involve lineage-specific parameters and only a subset of the genome (for instance, mature mRNAs only for most miRNAs), are a separate issue than determining the transcriptional origins.

We implemented alternative methods for handling MMAP sRNA-seq reads into version 3 of our general-purpose sRNA-seq analysis software, ShortStack (Axtell 2013), which is publicly available at https://github.com/MikeAxtell/ShortStack/releases. ShortStack takes in raw sRNA-seq data (in fasta, fastq, or color-space formats), and a corresponding reference genome, and performs alignment, annotation, and quantification of expressed small RNAs (Figure 1). In the first step of alignment (Figure 2A) ShortStack uses bowtie (Langmead *et al*. 2009) to identify all possible best-matched alignments for each read, subject to a default, user-adjustable limit of 50 alignments per read. ShortStack will then calculate a probability for each alignment according to one of three alternative methods. Consider an example read with two possible alignments to different loci in the genome (green-colored read in Figure 2B). Each of the two possible loci in this example have different "neighborhoods" of adjacent alignments of other reads. In random-placement mode, the probability of originating from a locus is simply 1/n, where n is the number of possible alignments; in the example read from Figure 2, each position thus has a probability of 50% (Figure 2C). This emulates the default behavior of bowtie (Langmead *et al*. 2009) and bwa (Li and Durbin 2009). In unique-weighting mode (U), the frequencies of uniquely aligned reads mapping within the vicinity of the alignment under consideration are tallied. For the example read in Figure 2, locus 1 has seven uniquely aligned reads in the vicinity, while locus 2 has only one (Figure 2D). Thus, in U mode, the probability of the example read originating from locus 1 is calculated as 87.5%, while the probability of originating from locus is calculated as 12.5% (Figure 2D). The fractional method (F) uses all reads mapping within the vicinity of an alignment. Unique reads in the vicinity lend full weight, while MMAP reads provide weights inversely proportional to their MMAP-value. For the example read in Figure 2, the weighted probability for originating from locus 1 is thus calculated at 73.7%, while the probability for originating from locus 2 is calculated as 26.3% (Figure 2E). The calculated probabilities are then used as weightings in a random number selection to designate the primary alignment for the read (Figure 2F), and all of the other possible alignments are marked as secondary alignments. Alternatively, ShortStack can be instructed to simply ignore (N) all MMAP reads (Figure 2G), emulating the behavior of bowtie with option ‐m 1 set, or that of Novoalign's default settings.

**Figure 1.**
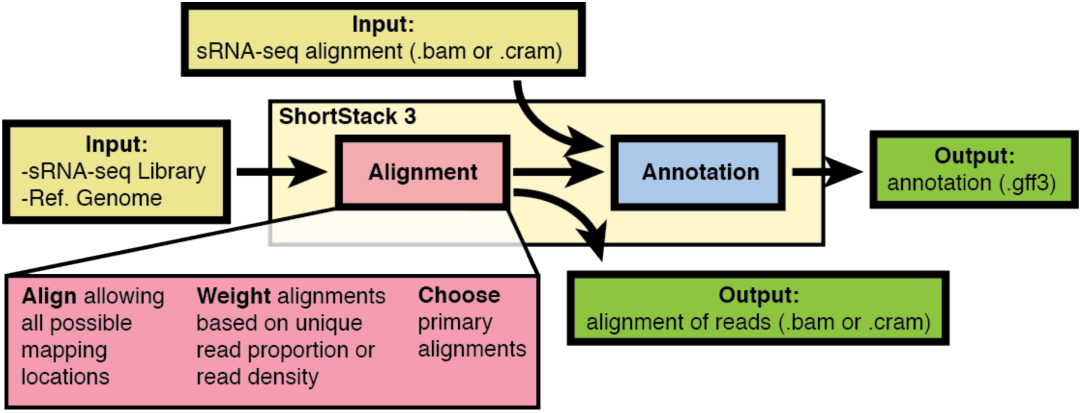
Overview of ShortStack methodology.

**Figure 2.**
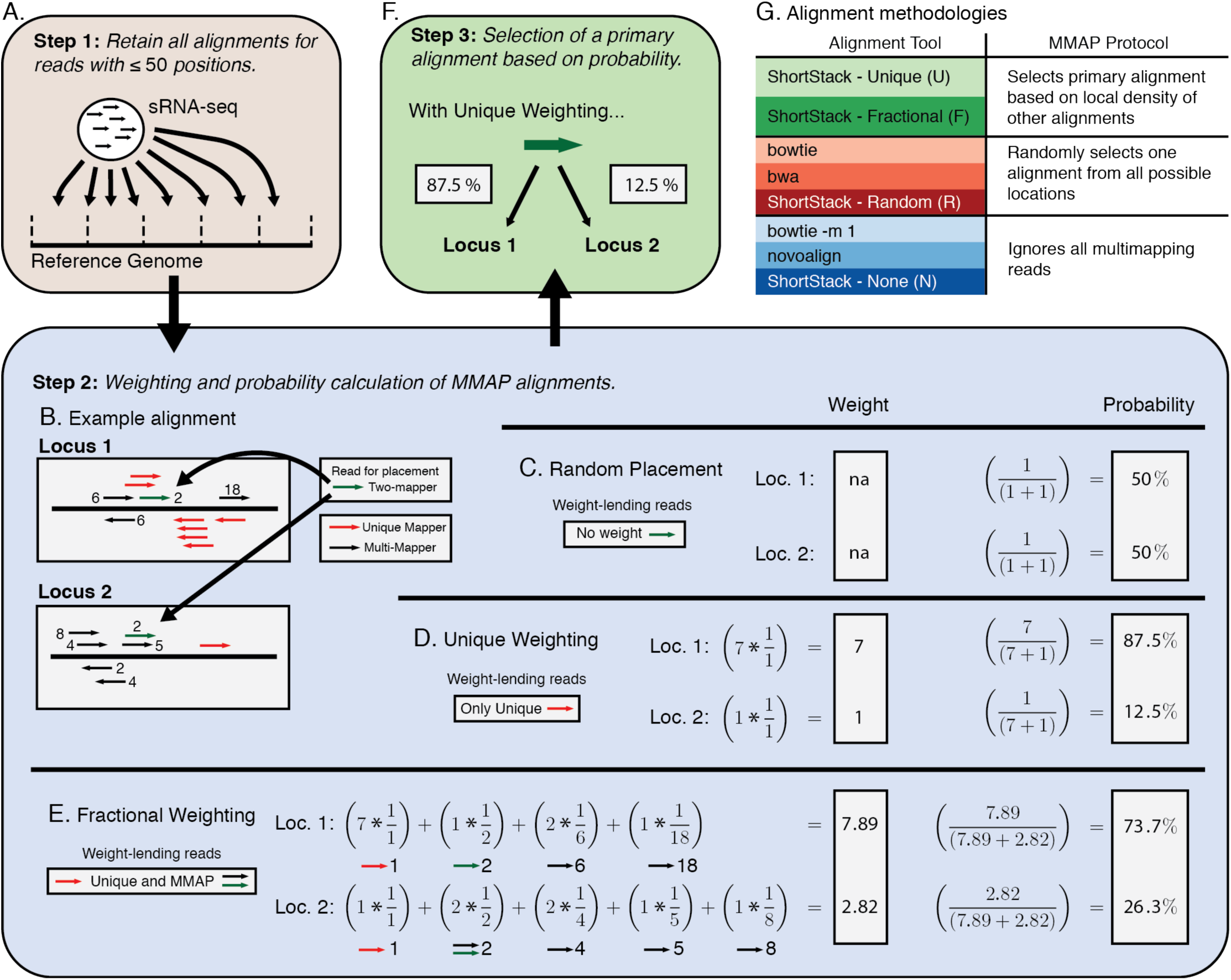
ShortStack3 alignment methodology. A) First step in alignment by ShortStack: Initial alignment of sRNA-seq reads to a reference genome. B) Example of local alignments for a read (green) with an MMAP-value of two. Numbers adjacent to reads indicate their MMAP-value. C) Weighting scheme for random placement of MMAP reads. D) Weighting scheme for ShortStack’s Unique method. E) Weighting scheme for ShortStack’s Fractional method. F) Final step: Choosing a single primary alignment based on calculated probabilities. 5 G) Alignment tools grouped by MMAP methods.

The vicinities used in methods U and F are obtained by first dividing the reference genome into 50 nt bins. The bin of a given alignment is defined as the location of its left-most aligned nucleotide. The vicinity is then defined as the total number of reads in a five-bin window, with the center bin being that of the alignment in question. Thus, the vicinity is essentially a 250 nt window centered on the alignment in question. When run in modes U or F, MMAP reads for which the computed probabilities at all positions are equal remain a random guess. In those cases, such reads will be suppressed entirely (marked as unmapped) if they had more than three possible alignment positions (this threshold is user-adjustable).

It's important to note that the U and F weighting methods are not 'winner-take-all'. For instance, if the U method was used for the example in Figure 2, there would be a 12.5% probability that the less-likely position would be marked as the primary alignment. Similarly, MMAP placement decisions are made on per-read basis, not a per-sequence basis. For example, 100 MMAP sRNA-seq reads of identical sequence with two possible alignment positions of probabilities 80% and 20%, we expect approximately 80 of the reads to be placed in the first position, and approximately 20 of them to be placed in the second position.

### Data Sources

sRNA-seq libraries from *Arabidopsis thaliana*, *Oryza sativa*, and *Zea mays* were obtained from the NCBI Sequence Read Archive (SRA) (Table S1). Libraries were selected that had over 5 million raw reads, were available in an unprocessed format, and were derived from an Illumina instrument. 3' adapter sequences were discovered using *fmd_3p_adapter.pl* (available at http://sites.psu.edu/axtell/) and removed using ShortStack's internal adapter trimming protocol. Simulated sRNA-seq libraries were produced to closely emulate real sRNA-seq data. This process was accomplished through a custom python script and wrapper run under default settings: *sRNA-simulator.py* (File S1). This script uses a real sRNA-seq library as the basis for each simulated library. Real sRNA-seq libraries were aligned using bowtie (Langmead *et al*. 2009) reporting all alignments. Regions of the genome which had no alignments were removed from consideration as simulated loci, while genomic regions prone to alignments with certain length classes of sRNAs became candidate regions for simulated heterochromatic siRNA (hc-siRNA; 23 - 24 nt) and trans-acting siRNA (21 nt) loci. miRNA candidate regions were picked based on prior annotated loci, available through miRBase (Kozomara and Griffiths-Jones 2014). Simulated loci were chosen from these candidate regions at random. Five million reads were then generated from these simulated loci, generating roughly 3.25M hc-siRNA, 1.5M miRNA and 250k tasiRNA reads. Loci were made to approximate real loci in size and pattern: hc-siRNA as primarily 24 nt RNAs from 200-1000 nt loci, from both genomic strands; miRNA as 21 nt RNAs from 125 nt loci with a miRNA and miRNA* pattern; tasiRNA as 21 nt RNAs from 140 nt loci producing a number of phased reads, from both genomic strands. All three loci types produced a realistic distribution of differently sized or shifted reads to simulate mis-processing. Sequencing errors are simulated at a rate of one mis-sequenced base per 10,000 reads. Unlike real data, simulated reads are traceable to their locus of origin, and thus are suitable to discern correct placements from incorrect ones. PolyA+ mRNA-seq data were obtained from SRA (Table S1). Reference genome versions were TAIR10 (*Arabidopsis thaliana*), IRGSP7 (*Oryza sativa*), and B73v3 (*Zea mays*).

### Alignments and analyses

sRNA-seq libraries were aligned using ShortStack (Axtell 2013), bowtie (Langmead *et al*. 2009), bwa (Li and Durbin 2009) and Novoalign (Novocraft.com). Specific versions and settings are specified in Table S2. mRNA-seq libraries were trimmed using cutadapt (Martin 2011) or processed as needed to de-pair paired-end datasets, and aligned using tophat2 (Kim *et al*. 2013) with bowtie (Langmead *et al*. 2009) as its alignment algorithm. Data processing details for each analysis are listed in Table S1.

### Analysis of Pol IV / RDR2-dependent siRNAs

Candidate RNA polymerase IV / RDR2 (P4R2)-dependent siRNA precursors were obtained from RNA-seq data from *Arabidopsis thaliana dcl234* triple mutants (Li *et al*. 2015; Blevins *et al*. 2015; Zhai *et al*. 2015; Ye *et al*. 2016). Alignment was performed using bowtie, tolerating no mismatches and retaining only uniquely aligned reads between 28 and 60 nts in length. These reads were then used to computationally generate four 24 nt siRNA ‘daughters’ from each putative precursor, corresponding to both ends of top and bottom strands of the presumed duplex. Computational daughters that were actually sequenced in the corresponding wild-type sRNA-seq libraries (Table S1) were noted. The wild-type sRNA-seq data were aligned using different alignment approaches. The precision of the tracked MMAP daughter reads was assessed using the coordinates of the corresponding uniquely aligned precursor as the known true origins.

### sRNA-seq

Total RNA was isolated from wild-type Col-0 *Arabidopsis thaliana* inflorescences using Tri-Reagent per the manufacturer’s instructions (ThermoFischer, 4368814). sRNA-seq libraries were constructed using the Tru-Seq sRNA kit (Illumina, RS-200-0012) per the manufacturer's instructions, and sequenced on an Illumina Hi-Seq 2500. sRNA-seq data has been deposited at NCBI GEO under accession GSE76281 (Table S1).

### qRT-PCR of primary miRNA transcripts

Total RNA was isolated from wild-type Col-0 *Arabidopsis thaliana* inflorescences using Tri-Reagent per the manufacturer’s instructions (Thermo Fisher, AM9738). Primers were designed using the NCBI PrimerBlast tool (Ye *et al*. 2012) and targeted the 3' region downstream of the Dicer-Like (DCL) cleavage sites. Primer sequences are given in Table S3. cDNA was synthesized by reverse transcription kit (Thermo Fisher, 4368814) and qRT-PCR experiments run on a Life Technologies StepOne Plus real-time PCR instrument using SYBR Green-based master mix (Quanta Biosciences, 95073-012) per manufacturer's instructions. Expression is calculated as the fluorescence relative to Actin (ACT2), calculated using grouped PCR efficiencies from analysis by LinRegPCR (Ramakers *et al*. 2003; čikoś *et al*. 2007).

### Data availability

Information on sequencing libraries generated in this study are available through NCBI GEO, accession number GSE76281. Information and accessions for all other libraries used are available in Table S1. The sRNA-seq simulator script is provided as File S1 and all other scripts used in analysis are available upon request. Sequences and annotation information for miRNAs were acquired through miRBase (Kozomara and Griffiths-Jones 2014).

## Results and Discussion

### Prevalence of MMAP reads in sRNA-seq data

In a survey of polyA+ mRNA-seq data from three plant species, MMAP percentages above 10% are rare (Figure 3A). In contrast, sRNA-seq datasets from the same three species frequently have over 50% MMAP reads (Figure 3A). This illustrates the extent of the MMAP problem specifically in sRNA-seq data. We examined a collection of studies (n = 20) where plant sRNA-seq data were aligned to a reference genome (Figure 3B, Table S4). In 80% of the cases, MMAP reads were handled by random selection, where a single possible alignment is randomly selected for each MMAP read. Although computationally easy, random selection is highly imprecise, as most random choices will be wrong. Another 10% of the studies examined simply ignored MMAP reads. This method has the clear disadvantage of discarding huge amounts of data, often over 50% (Figure 3A). This survey of data and published studies clearly indicates that MMAP reads are a large issue in sRNA-seq data analysis that are often dealt with ineffectively.

**Figure 3.**
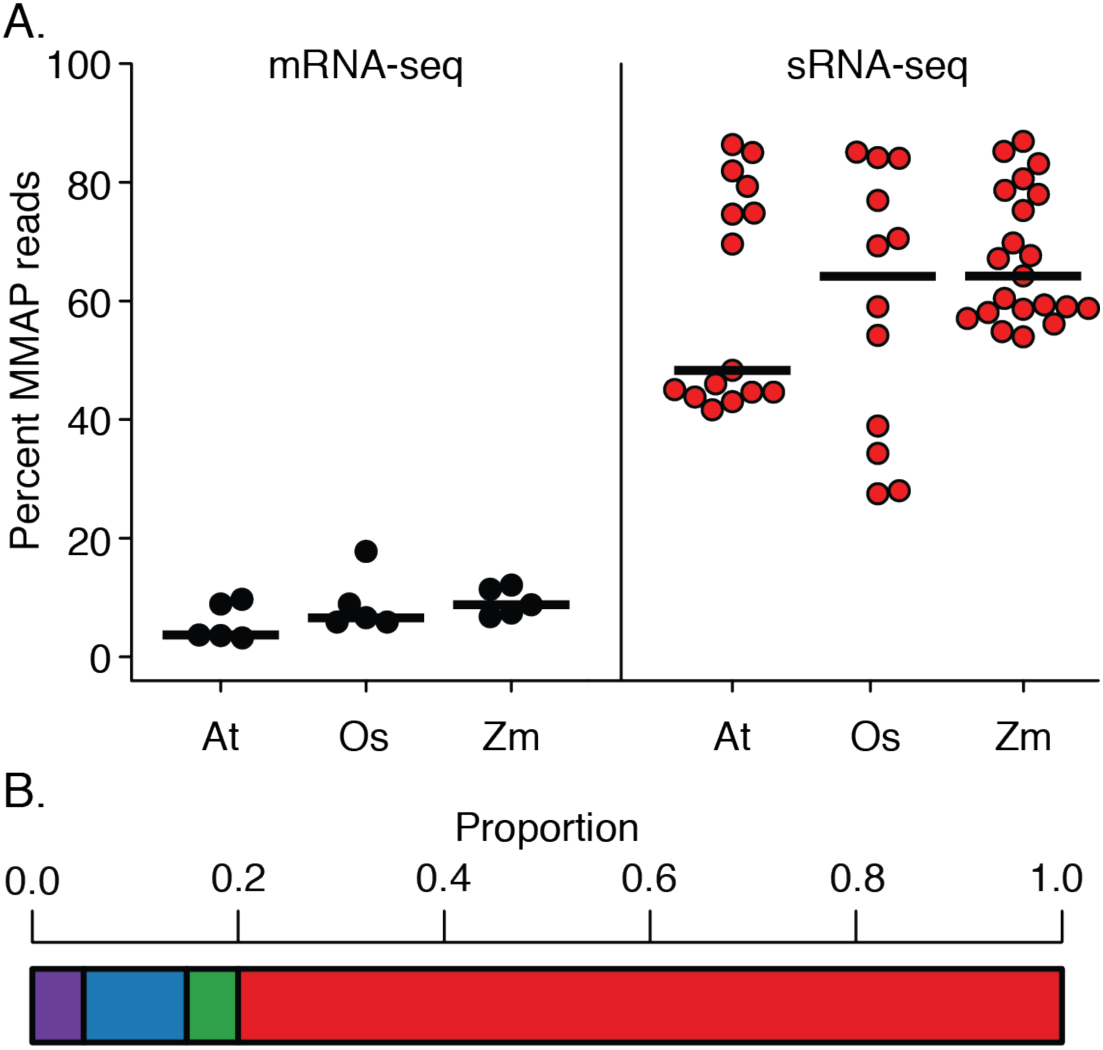
Prevalence of MMAP reads and methods A) MMAP rates for reads from mRNA-seq and sRNA-seq data from *Arabidopsis thaliana* (At), *Oryza sativa* (Os), and *Zea mays* (Zm). Horizontal bars mark median values, circles mark values for individual libraries. B) Proportion of MMAP-selection methods for sRNA alignment in recent literature (n = 20; Table S4).

### High-precision alignments of MMAP sRNA-seq reads

We designed two alternative modes of proximity-based weighting systems (U and F) to place MMAP reads, and implemented them into the sRNA-seq analysis software ShortStack (Figure 2) (Axtell 2013). We then conducted a performance analysis comparing these methods to several other aligners with various settings. The alignment methods can be classified into three groups based on their treatment of MMAP reads (Figure 2): ShortStack-U and ‐F both use a proximity-weighted scheme to influence placement of MMAP reads. In contrast, under their default settings bowtie (Langmead *et al*. 2009) and bwa (Li and Durbin 2009) randomly place MMAP reads, as will ShortStack when run in the R mode. Finally, Novoalign, bowtie with setting ‐m 1, and ShortStack in mode N all will simply ignore MMAP reads. We simulated sRNA-seq data based on the properties of real datasets from *Arabidopsis thaliana*, *Oryza sativa*, and *Zea mays*. Unlike real data, the ‘correct’ alignment location for each MMAP is known for simulated data. This allows direct comparison of aligner performance. Specifically, we can count true positives (aligned reads that are put in the correct position; TP), false positives (aligned reads that are put in the incorrect position; FP), true negatives (unaligned reads that have no possible alignment position; TN), and false negatives (unaligned reads where there is at least one valid alignment position; FN). Precision (TP/(TP + FP)) and sensitivity (TP/(TP + FN)) can then be calculated in the standard manner.

In nearly all cases, the highest precisions were obtained with ShortStack’s U method (Figure 4A).

**Figure 4.**
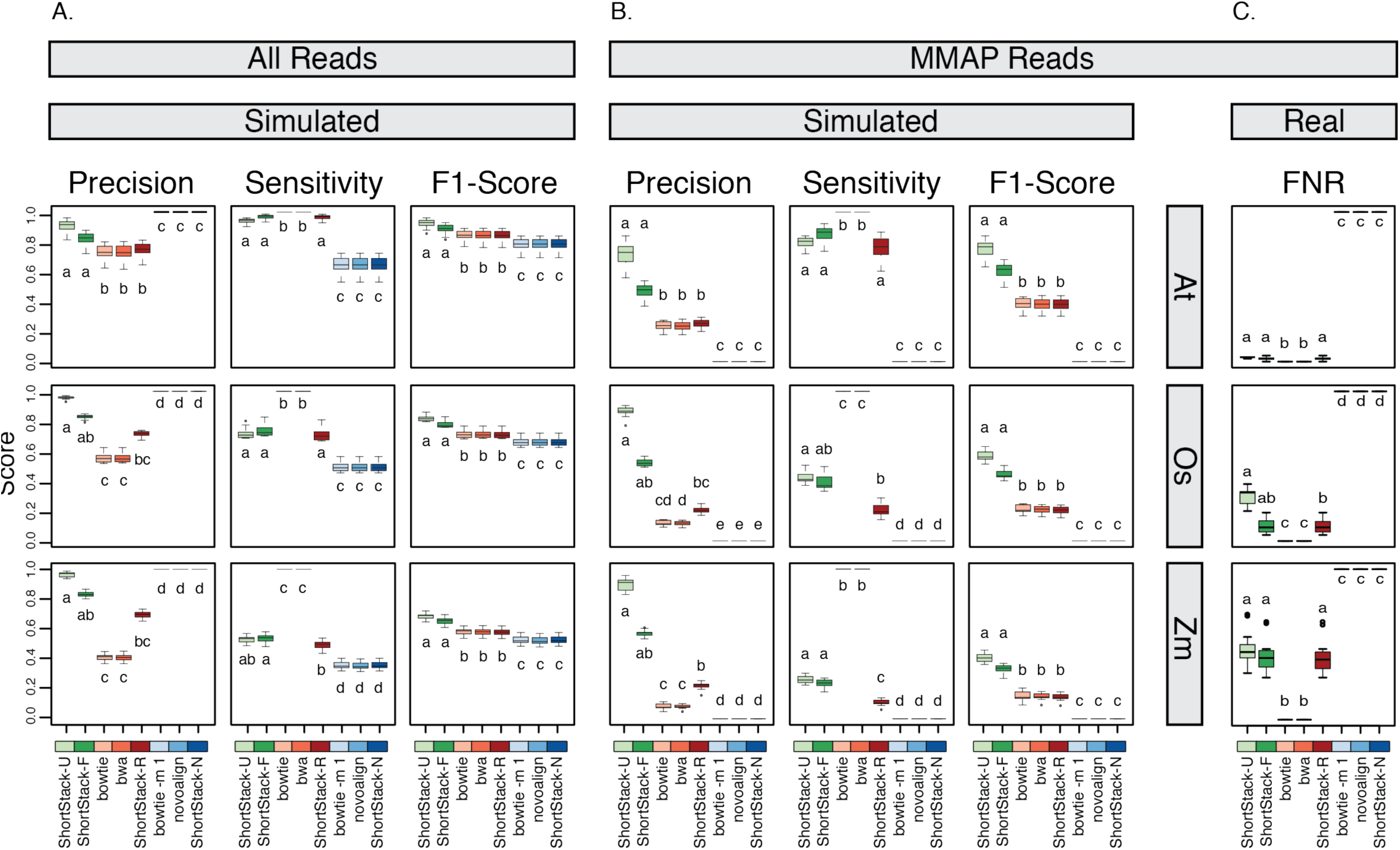
Performance analysis of sRNA-seq alignment methods. A) Precisions, sensitivities, and F1 scores for alignments of simulated sRNA-seq data with the indicated methods for entire datasets. Boxplots show medians (central bars), the 1st to 3rd quartile range (boxes), other data out to 1.5 the interquartile range (whiskers), and outliers (dots). n=15, 12, and 21 for the At, Os, and Zm data, respectively. Treatments sharing a common letter indicate groups that are not significantly different by non-parametric analysis (Kruskal–Wallis ANOVA with Dunn multiple comparison test, α = 0.05). At: *Arabidopsis thaliana*, Os: *Oryza sativa, Zm: Zea mays*. B) MMAP reads only. Same analysis and plotting conventions as in panel A. C) False negative rates for alignments of real sRNA-seq data with the indicated methods for MMAP reads. Plotting conventions as in panel A.

ShortStack-U had significantly (Kruskal–Wallis ANOVA with Dunn multiple comparison test, α = 0.05) higher precisions than any non-ShortStack method. The results were especially striking when examining the precision specifically for MMAP reads. ShortStack’s U method routinely achieved precision values of 75% or higher (Figure 4A). In contrast, none of the random-placement or ignore MMAP methods ever had MMAP precision values over 50%. The high precision does come at the cost of sensitivity: As described in the methods, ShortStack-U and ‐F will not place all MMAP reads. Instead, if a read has too many possible positions (default is > 50) or if the weighted probabilities are equal, MMAP reads are suppressed. Depressed sensitivities are more apparent in the more repetitive genome of *Zea mays* as compared to *Arabidopsis thaliana* (Figure 4A). We computed F1 scores (harmonic means of precision and sensitivity) to assess the balance between precision and sensitivity. By this metric, ShortStack-U and ‐F both performed significantly (Kruskal–Wallis ANOVA with Dunn multiple comparison test, α = 0.05) better than any other method in all three species (Figure 4A).

Analysis of real libraries is limited to examination of false negative rates (FNR = FN / (FP + TP)), as TP and FP cannot be distinguished. As expected, FNRs are higher in *Zea mays* and *Oryza sativa* relative to *Arabidopsis thaliana*, reflecting the higher proportions of reads with high MMAP-values from the more repetitive genomes (Figures 4C). Under default settings, ShortStack’s U and F methods discard ~30-50% of the MMAP reads from rice and maize, but are likely placing the remainder with very high confidence.

### Precision is a function of MMAP-value

As discussed above, our methods by default will not attempt weighting or placement of MMAP reads with more than 50 possible alignment positions. This is based on the prediction that for reads with very high MMAP-values, we will have an unacceptably high error rate. We tested this prediction by examining the precision values in our simulated datasets as a function of MMAP value (Figures 5A–B). As expected, precision declines as a function of higher MMAP values in all cases (Figure 5A), with random having the most severe drop-off. Notably, the U method consistently gives the highest precisions at a given MMAP value. Analysis of cumulative precisions showed that plateaus of precision generally became apparent at MMAP-values ~ 50 (Figure 5B), justifying our default settings to ignore reads with more than 50 possible alignment positions. Our simulated data had more high MMAP reads than real data (Figure 5C), indicating that aligner performance with real data is likely to be superior to that of estimated using simulated reads.

**Figure 5.**
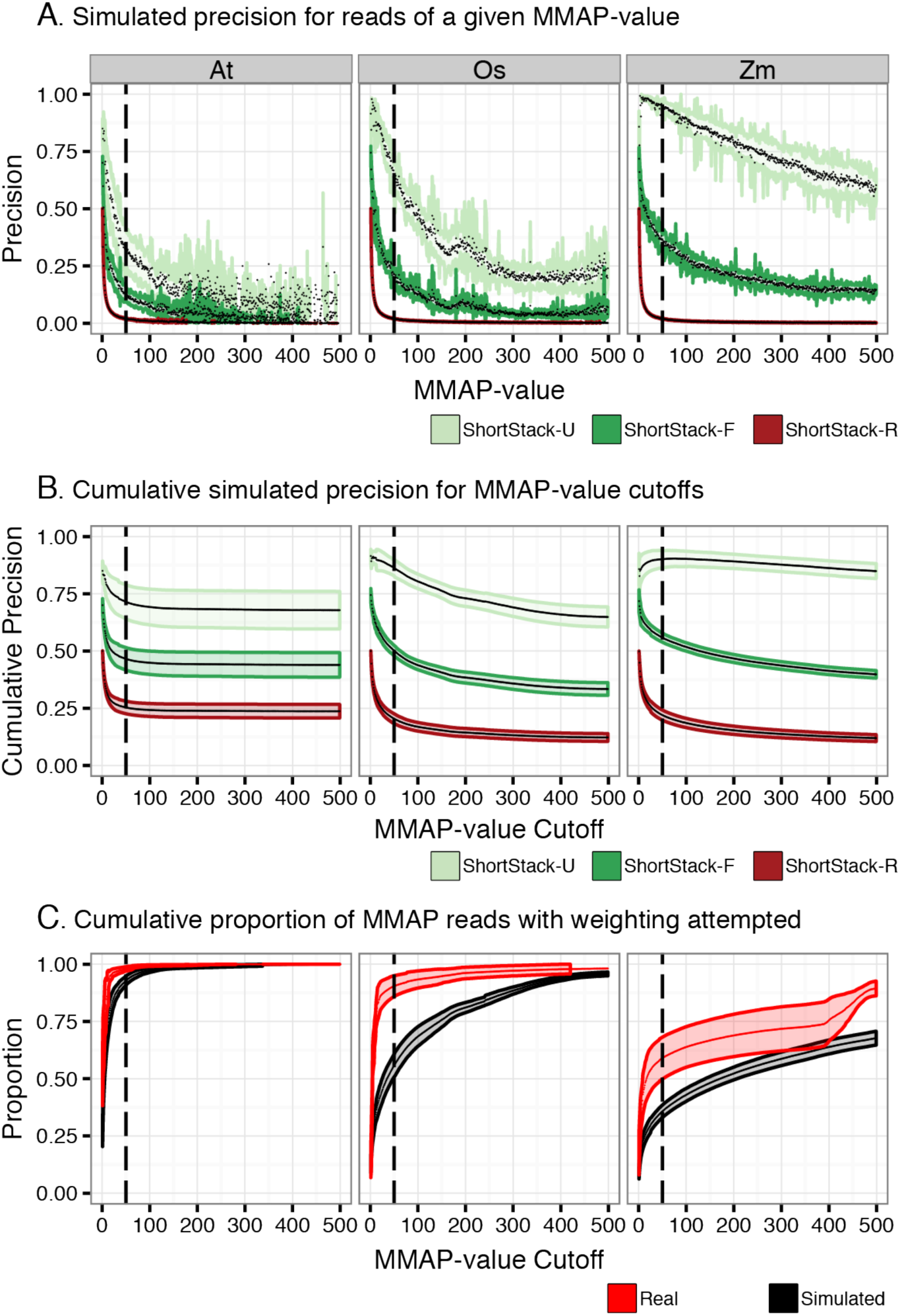
Influence of MMAP-value on performance A) Precision as a function of MMAP-value for simulated sRNA-seq data from the indicated species and alignment method. MMAP-value is the number of possible alignment positions for a read. Colored lines are standard deviations, black dots are mean values. Heavy dashed line at MMAP=50 indicates the default cutoff value for ShortStack, above which placement of MMAP reads is not attempted. At: *Arabidopsis thaliana*, Os: *Oryza sativa*, Zm: *Zea mays*. B) Cumulative precision as a function of MMAP-value for simulated sRNA-seq data from the indicated species and alignment method. Plotting conventions as in panel A. C) Cumulative proportion of real and simulated sRNA-seq data retained by ShortStack alignments under differing MMAP-value cutoffs. Note that simulated libraries have higher proportions of reads with high MMAP values. Plotting conventions as in panel A.

### Experimental testing of alignment methods

Mature miRNAs are frequently encoded by multiple paralogous loci, which causes sRNA-seq reads corresponding to the mature miRNA to be MMAPed. However, the primary transcripts derived from each of the paralogs typically are much longer and have distinct sequences, such that qRT-PCR approaches can distinguish them. We used accumulation levels of primary transcripts as proxies for the true expression levels of various *Arabidopsis thaliana MIRNA* loci that encode identical mature miRNAs, and compared those data to estimates derived from small RNA-seq alignments (Figure 6). qRT-PCR indicated sharp differences in primary transcript accumulation for each of the three miRNA families we studied (Figure 6A). ShortStack's U method produced mature miRNA alignments that were mostly similar to the results from qRT-PCR of the primary transcripts (Figure 6B). In contrast, the other alignment methods suggested nearly equal accumulation of mature miRNAs from each of the paralogs. We quantified the fit between the qRT-PCR data and the sRNA-seq alignment data by analyzing the squared residual errors between the scaled expression levels (Figure 6C). By this metric, ShortStack-U predicts the specific contributions of miRNA paralogs significantly (Kruskal–Wallis ANOVA with Dunn multiple comparison test, α = 0.05) better than all other tested methods.

**Figure 6.**
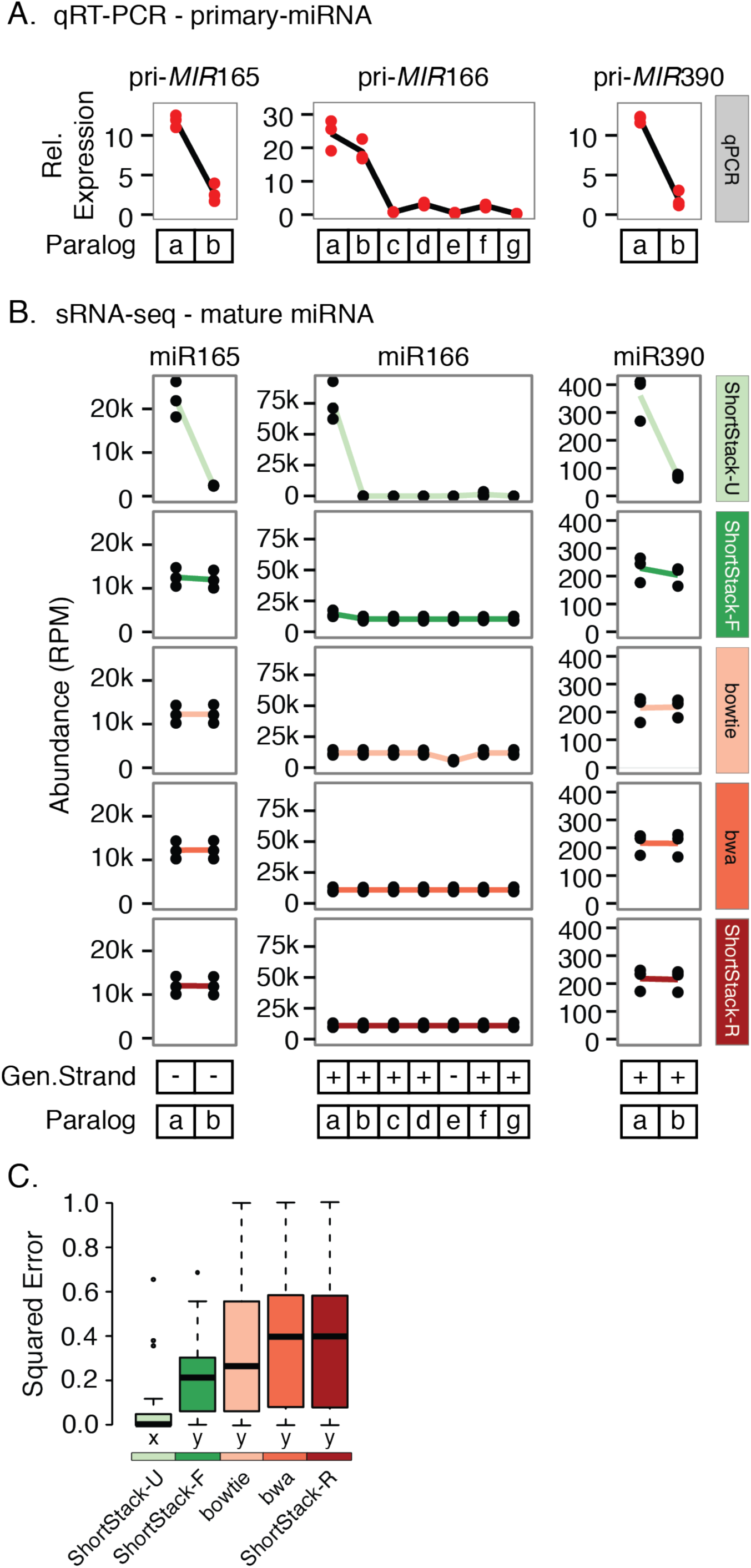
Experimental assessment of sRNA-seq alignment methods using miRNA paralogs A) Relative expression of the indicated primary *MIRNA* transcripts in *Arabidopsis thaliana* Col-0 inflorescences assessed via qRT-PCR. Values are normalized to 1/1000 those of *ACTIN2*. Dots show values from biological replicates (n=3). B) Accumulation of the indicated mature miRNAs from each of their possible paralogs as determined by different sRNA-seq alignment methods. Values are from three biological replicate sRNA-seq libraries from *Arabidopsis thaliana* Col-0 inflorescences. C) Squared residual errors from comparisons of scaled qRT-PCR data to scaled sRNA-seq alignment results. Boxplots show medians (horizontal bars), the 1st to 3rd quartile range (boxes), data out to 1.5 times the inter-quartile range (whiskers), and outliers (dots). Treatments sharing a common letter indicate groups that are not significantly different by non-parametric analysis (Kruskal–Wallis ANOVA with Dunn multiple comparison test, α = 0.05).

When DCL enzymes are absent from *Arabidopsis thaliana*, longer double-stranded RNAs accumulate that are dependent on RNA polymerase IV (Pol IV) and RNA-dependent RNA polymerase 2 (RDR2) (Li *et al*. 2015; Blevins *et al*. 2015; Zhai *et al*. 2015; Yang *et al*. 2016; Ye *et al*. 2016). These are likely to be the direct substrates for the DCL3-mediated production of 24 nt siRNAs by excision from the ends of precursors. We identified a set of longer precursors from *dcl2/dcl3/dcl4* triple mutant plants that could be uniquely mapped to the reference genome. We then computationally diced this set of precursors to create predicted mature 24 nt siRNAs, and examined the corresponding wild-type sRNA-seq libraries to find cases where the putative mature siRNAs actually were sequenced. The wild-type sRNA-seq libraries were aligned with various methods. Cases where the mature siRNAs of interest were MMAP allowed assessment of alignment precision, as we presume their uniquely aligned precursors represent the correct alignment positions. ShortStack-U significantly (Kruskal–Wallis ANOVA with Dunn multiple comparison test, α = 0.05) outperformed other alignment methods (Figure 7) in this analysis, as measured by precision. Thus, experimental data from both miRNAs and endogenous siRNAs supports the hypothesis that ShortStack's U alignment method is superior.

**Figure 7.**
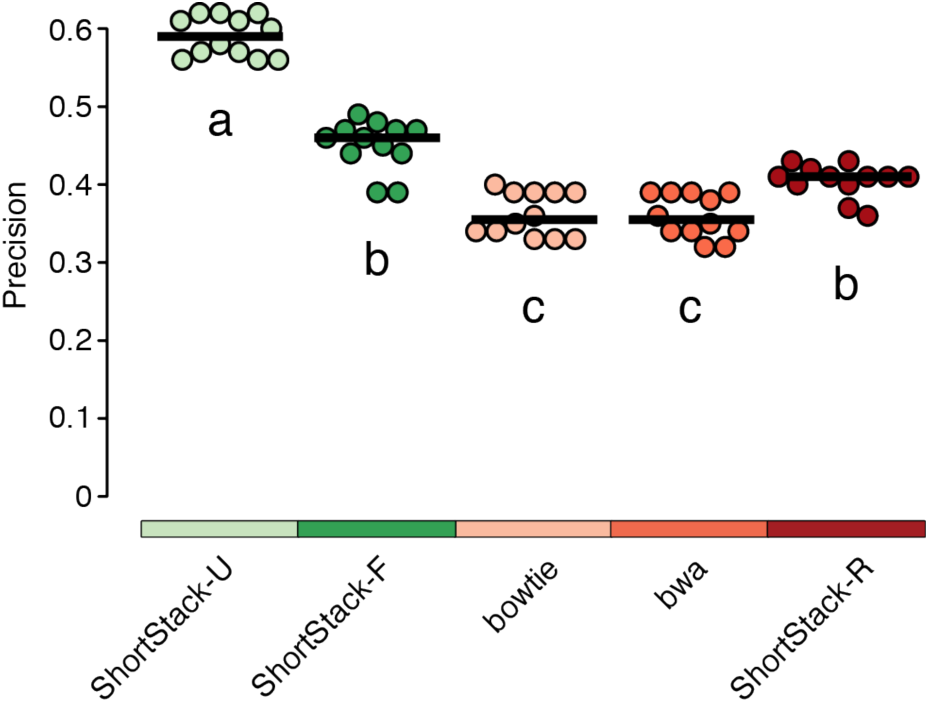
Precisions from alignments of *Arabidopsis thaliana* MMAP 24 nt siRNAs whose true origins are known based on a unique precursor alignment. Dots: data from individual libraries; horizontal bars: medians. Treatments sharing a common letter indicate groups that are not significantly different by non-parametric analysis (Kruskal–Wallis ANOVA with Dunn multiple comparison test, α = 0.05).

### Bowtie shows bias in random read placement

When run under default settings, bowtie (Langmead *et al*. 2009) randomly selects one possible alignment position for MMAP reads. However, the software documentation warns that biased strand selection, where alignment positions from one strand are favored, can occur (Langmead *et al*. 2009). For sRNA-seq data, we have observed that this strand bias is quite strong. We observe a strong top strand bias when bowtie is run under default settings to align sRNA-seq data (Figure 8). Indeed, this can affect estimation of miRNA accumulation from paralogs (Figure 6B, miR166e). This bias is not apparent when other methods are used (Figure 8). Small RNA polarity is an important feature used for annotations, so it is conceivable that this strand bias could distort results.

**Figure 8.**
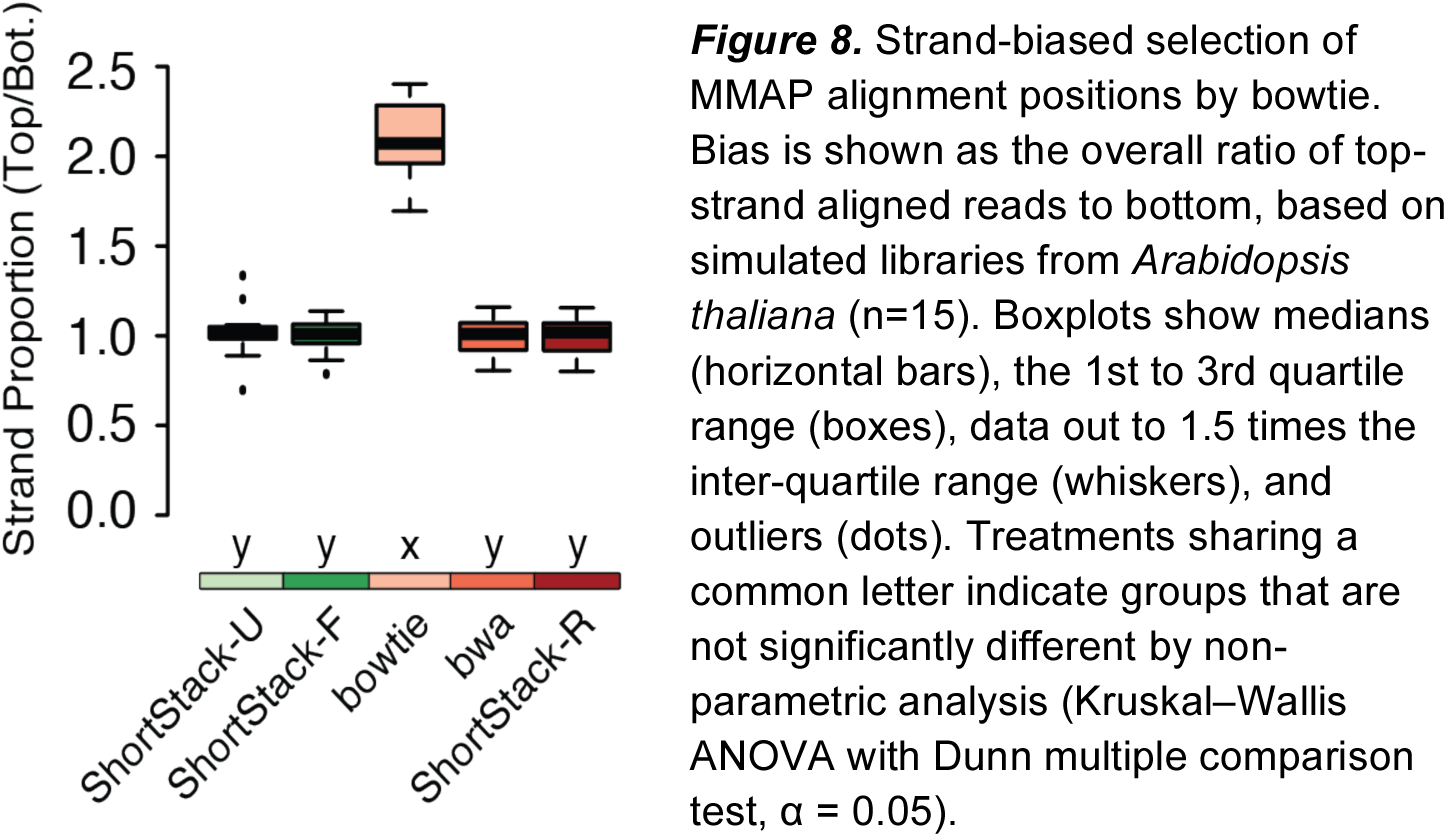
Strand-biased selection of MMAP alignment positions by bowtie. Bias is shown as the overall ratio of top-strand aligned reads to bottom, based on simulated libraries from *Arabidopsis thaliana* (n=15). Boxplots show medians (horizontal bars), the 1st to 3rd quartile range (boxes), data out to 1.5 times the inter-quartile range (whiskers), and outliers (dots). Treatments sharing a common letter indicate groups that are not significantly different by nonparametric analysis (Kruskal–Wallis ANOVA with Dunn multiple comparison test, α = 0.05)

### Speed testing

We compared the alignment speed of our methods with those of other aligners. To account for differences caused by different genome sizes and/or sRNA-seq libraries, we measured speed in normalized units. Because bowtie under default settings was always the fastest method, we normalized each run to a multiple of the bowtie-default completion speed. The time spent on genome-indexing was not part of this assessment. In general, our favored method, ShortStack-U, was five to ten times slower than bowtie-default, but comparable to bwa (Figure 9). Given that the results of ShortStack-U are superior to the other methods, we feel that the extra time is an acceptable trade-off. In absolute terms, the wall-time for our ShortStack-U runs on these data ranged from 6 minutes to 2.5 hours, compared with bowtie-default which ranged from 1 to 32 minutes.

**Figure 9.**
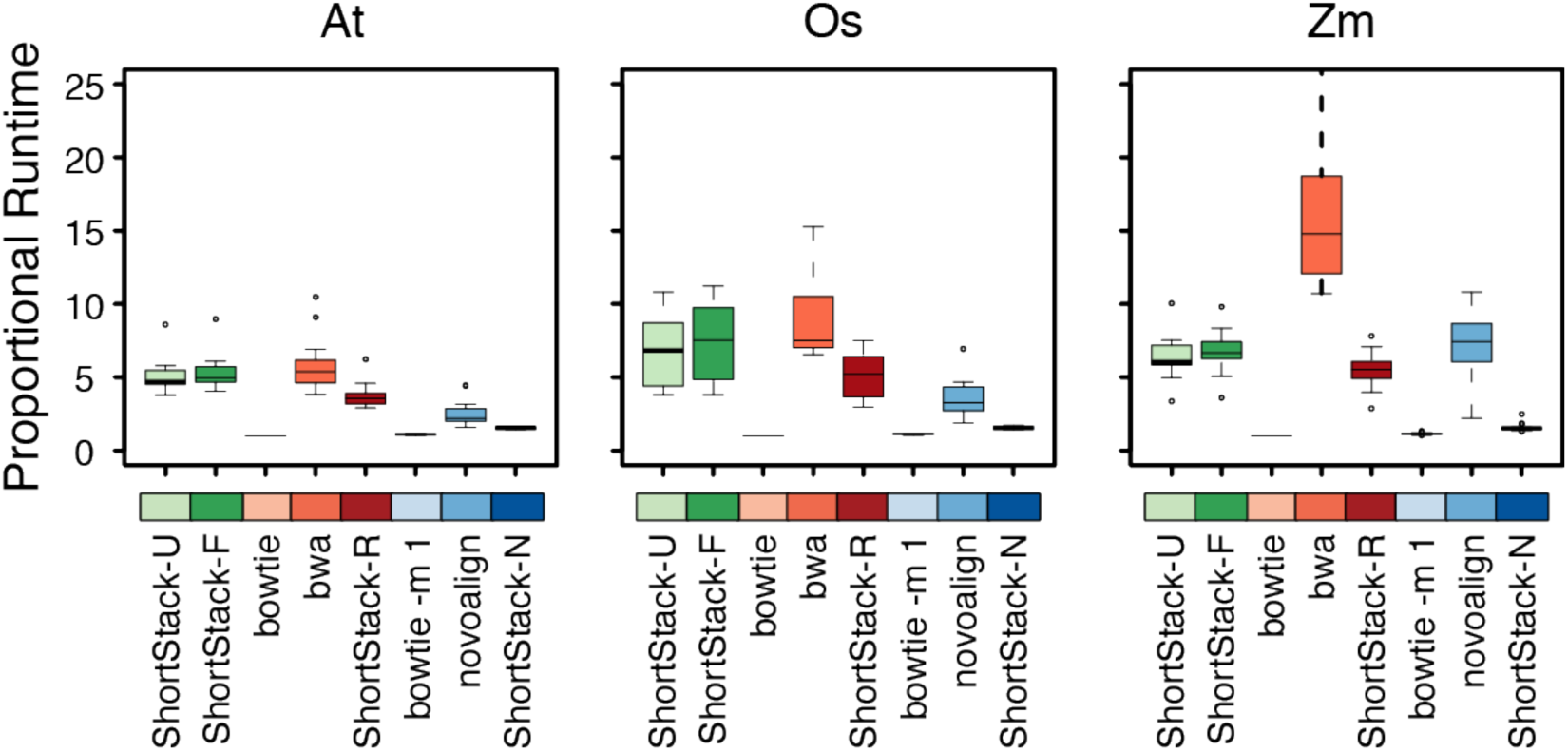
Comparison of alignment times for real sRNA-seq libraries with the indicated methods. Boxplots show medians (central bars), the 1st to 3rd quartile range (boxes), other data out to 1.5 the interquartile range (whiskers), and outliers (dots). n=15, 12, and 21 for the At, Os, and Zm data, respectively. At: *Arabidopsis thaliana*, Os: *Oryza sativa*, Zm: *Zea mays*.

## Conclusions

A particularly important goal of ours is to reduce the likelihood of false discoveries in small RNA gene annotation. Reducing the number of incorrect alignments is likely to reduce spurious annotations. ShortStack-U gives fewer false positives for MMAP reads than all other methods for sRNA-seq alignment (Figure 4, 5, 7). The experimental evidence identifying miRNA alignment is striking, as mature miRNA are remarkably poorly placed through standard approaches (Figure 6). It is likely that misalignment of this nature is significantly misrepresenting the abundance of many loci, conceivably resulting in mis-annotations. ShortStack-U's alignment method reduces these risks, striking a balance between precision and sensitivity.

A method which is capable of identifying the true site of origin for all MMAP sRNA-seq reads is elusive. To achieve reasonable precision, our methods sacrifice some sensitivity by ignoring highly MMAP reads. However, it should be kept in mind that these reads are part of the small RNA profile too and should not be completely ignored. Future development of sRNA-seq alignment methods should focus on increasing precisions to allow confident placement of even the most repetitive reads. Until then, these most highly-repetitive reads can be analyzed using methods independent of alignment to the reference genome.

Overall, our work demonstrates that sRNA-seq alignments can be significantly improved by using local weighting to guide placement of multi-mapped reads, and that biases in placement of multi-mapped reads can influence downstream analyses.

## Author Contributions

MJA developed ShortStack code. NRJ and MJA developed alignment algorithm ideas and ideas for performance testing. NRJ conducted performance analyses and generated all figures. NRJ and JY conducted qRT-PCR experiments. CC constructed sRNA-seq libraries. MJA and NRJ wrote the manuscript with comments from JY and CC.

## Acknowledgements

This work was supported by an award from the US NSF to MJA (award # 1339207). sRNA-seq was conducted on an Illumina HiSeq2500 purchased with funds from a US NSF award to MJA (award #1229046)

## Supplemental Data

Supplemental Table 1: Dataset accession numbers and descriptions

Supplemental Table 2: Versions and settings

Supplemental Table 3: Oligo sequences

Supplemental Table 4: Methods used for placement of MMAP sRNA-seq reads in 20 previous studies.

Supplemental File 1: *sRNA-simulator.py*: Python script used to create simulated sRNA-seq datasets.

